# Replication Protein A1 is essential for DNA damage repair during mammalian oogenesis

**DOI:** 10.1101/2023.07.04.547725

**Authors:** Xiaosu Miao, Rui Guo, Andrea Williams, Catherine Lee, Jun Ma, P. Jeremy Wang, Wei Cui

## Abstract

Persistence of unrepaired DNA damage in oocytes is detrimental and may cause genetic aberrations, miscarriage, and infertility. RPA, an ssDNA-binding complex, is essential for various DNA-related processes. Here we report that RPA plays a novel role in DNA damage repair during postnatal oocyte development after meiotic recombination. To investigate the role of RPA during oogenesis, we inactivated RPA1 (replication protein A1), the largest subunit of the heterotrimeric RPA complex, specifically in oocytes using two germline-specific *Cre* drivers (*Ddx4-Cre* and *Zp3-Cre*). We find that depletion of RPA1 leads to the disassembly of the RPA complex, as evidenced by the absence of RPA2 and RPA3 in RPA1-deficient oocytes. Strikingly, severe DNA damage occurs in RPA1-deficient GV-stage oocytes. Loss of RPA in oocytes triggered the canonical DNA damage response mechanisms and pathways, such as activation of ATM, ATR, DNA-PK, and p53. In addition, the RPA deficiency causes chromosome misalignment at metaphase I and metaphase II stages of oocytes, which is consistent with altered transcript levels of genes involved in cytoskeleton organization in RPA1-deficient oocytes. Absence of the RPA complex in oocytes severely impairs folliculogenesis and leads to a significant reduction in oocyte number and female infertility. Our results demonstrate that RPA plays an unexpected role in DNA damage repair during mammalian folliculogenesis.

## Introduction

As the largest cell in the body of mammals, the oocyte is crucial for female fertility and early embryogenesis. To acquire maturity and competency for fertilization and subsequent embryonic development, female germ cells undergo a series of developmental processes. In mammals, following premeiotic DNA replication, oocytes enter an extended prophase of meiosis I during fetal development. In mice, oocytes progress through leptotene and zygotene stages, and reach the pachytene stage by embryonic day 18.5 (E18.5) (Borum, 1961; Pepling, 2006). At postnatal day 5 (P5), the majority of oocytes reach and arrest at the late diplotene (or dictyate) stage, with individual oocytes contained in the primordial follicles where they remain arrested until ovulation (Cohen et al., 2006; Pepling and Spradling, 2001). In mice, cohorts of primordial follicles are periodically recruited to enter a 3-week growth phase, progressing through primary and secondary follicles (Amleh and Dean, 2002). After puberty, these pre-antral follicles are further stimulated by follicle-stimulating hormone (FSH), leading to the formation of antral follicles (Barnett et al., 2006; Kumar et al., 1997). Throughout these stages, oocytes remain arrested at the diplotene or germinal vesicle (GV) stage due to their interactions with the surrounding granulosa cells (Coticchio et al., 2015). On the other hand, oocytes also control ovarian follicular development and the proliferation of granulosa cells (Eppig, 2001). Finally, triggered by the luteinizing hormone (LH) surge, fully grown immature oocytes are released from the GV stage arrest, resume meiosis, and progress through metaphase I (MI) and metaphase II (MII), leading to ovulation (Richards et al., 2002). Any disruption during meiotic divisions can lead to improper segregation of chromosomes which can cause severe birth defects in the resulting progeny (Biswas et al., 2021; Sun and Cohen, 2013).

Because the prolonged arrest of mammalian oocytes at the diplotene or GV stage can last for months in mice and even decades in humans, there is a potential for the accumulation of DNA damage during this extended period (Carroll and Marangos, 2013). Alongside single-strand breaks (SSBs) and double-strand breaks (DSBs), other types of DNA lesions, such as base mismatches and alterations in DNA structure, can also disrupt DNA function during replication and transcription. Consequently, cells actively engage in DNA damage response and DNA damage checkpoints to ensure that damage is repaired before cell cycle progression occurs, or alternatively, to induce apoptosis-mediated cell death if the damage is severe and irreparable (Pailas et al., 2022; Rinaldi et al., 2020). Although the GV stage of oocytes is equivalent to the G2/M transition of somatic cells, several differences are known: 1) In primordial follicles, it is trans-activating p63 (TAp63), but not p53, that is activated after DNA damage induction, causing apoptosis of the primordial follicle stage oocytes (Luan et al., 2022; Suh et al., 2006); 2) TAp63 expression diminishes as the follicles and oocytes grow, with its expression absent in antral follicles (Livera et al., 2008; Suh et al., 2006); 3) Fully grown GV oocytes in antral follicles exhibit a weakened G2/M checkpoint, allowing oocytes with DNA damage to still progress into the M phase due to a limited ability to activate the ataxia telangiectasia mutated (ATM) kinase (Marangos and Carroll, 2012) and/or a delayed response to the DNA damage (Subramanian et al., 2020); 4) Oocytes with DNA damage that enter the M phase normally arrest at MI due to the activation of the spindle assembly checkpoint (SAC) (Collins et al., 2015).

During DNA replication and transcription, the double-stranded DNA (dsDNA) must be unwound and separated to act as a template for DNA or RNA synthesis. This unwinding exposes single-stranded DNA (ssDNA) regions, which are more susceptible to DNA damage factors and can accumulate DNA lesions (Saini and Gordenin, 2020). To safeguard and stabilize the exposed ssDNA, cells employ various ssDNA protecting complexes. Among these, the replication protein A (RPA) complex, composed of RPA1, RPA2, and RPA3, is the primary ssDNA-binding heterotrimeric complex involved in numerous DNA metabolic pathways, including DNA replication, recombination, repair, and DNA damage checkpoints (Wold, 1997; Zou et al., 2006). RPA binds ssDNA with high affinity, thereby preventing the formation of secondary structures and protecting ssDNA from the action of nucleases. During these processes, RPA also directly interacts with other DNA processing proteins (Deng et al., 2015). For instance, during homology-directed repair of DSBs, ssDNA tails are initially bound by RPA, which is then replaced by RAD51 recombinase (Stauffer and Chazin, 2004). During DNA damage checkpoints, RPA is required for localization of ATR (ataxia telangiectasia mutated and Rad3-related) kinase to the DNA damage sites and for activation of ATR-mediated phosphorylation of the downstream targets (Zou and Elledge, 2003). In addition to its role in somatic cells, RPA functions in meiotic recombination in germ cells (Shi et al., 2019). Interestingly, MEIOB is a meiosis-specific paralogue of RPA1, the largest subunit of RPA (Luo et al., 2013; Souquet et al., 2013). MEIOB forms a dimer with SPATA22 and the MEIOB-SPATA22 dimer interacts with the RPA complex (Guo et al., 2020; Ribeiro et al., 2021; Xu et al., 2017). MEIOB binds to ssDNA and is essential for meiotic recombination in both males and females (Luo et al., 2013; Souquet et al., 2013).

While RPA has been investigated in yeast and cell lines, its role *in vivo* during mammalian development remains largely unexplored due to the early embryonic lethality resulting from *Rpa1* mutation in mice (Wang et al., 2005). Although RPA has been demonstrated to be crucial for meiotic recombination in mouse spermatogenesis (Shi et al., 2019), its role in female germ cell development has yet to be elucidated. In this study, we generated conditional knockouts of *Rpa1* in oocytes using two distinct *Cre* drivers (*Zp3-Cre* and *Ddx4-Cre*). Surprisingly, we discovered that RPA plays an essential role in DNA damage repair after meiotic recombination and is required for follicular development during oogenesis.

## Materials and Methods

### Ethics statement

All animal related procedures and methods were carried out in accordance with the approved guidelines and regulations. All animal experimental protocols were approved by the Institutional Care and Use Committees of the University of Pennsylvania and the University of Massachusetts Amherst.

### Generation of *Rpa1* oocyte-specific conditional knockouts (OcKOs)

Sperm carrying the *Rpa1* tm1a allele (Fig. 1A, C57BL/6N-*A^tm1Brd^ Rpa1^tm1a(KOMP)Wtsi^*/JMmucd, from MMRRC, UC-Davis) were used to fertilize the cumulus-oocyte-complexes (COCs) collected from superovulated C57BL/6N females (#005304, Jackson Laboratory) in the HTF medium (MR-070-D) supplemented with L-Glutathione reduced (GSH) and high Ca^2+^ as indicated in the MMRRC protocol. Fertilized oocytes with pronuclei were cultured in KSOM (MR106D) for 3 days to reach morula and early blastocyst stages, which were then transferred into the recipients via non-surgical embryo transfer in the Animal Models Core Facility at the Institute for Applied Life Sciences (IALS) (University of Massachusetts-Amherst). *Rpa1*^tm1a^ mice were genotyped by PCR (primers flanking FRT-left: F1 gaggacccacagtgaaagga and R1 ccacaacgggttcttctgtt; primers flanking FRT-right: F2 tctcatgctggagttcttcg and R2 attatgcacgtgccacagg) and confirmed by Sanger sequencing. *Rpa1*^tm1a^ mice were bred with FLPo (Raymond and Soriano, 2007) deleter mice where FLP recombinase is ubiquitously expressed (JAX, # 012930). Both germline transmission and successful FLP/FRT recombination were confirmed in the offspring carrying the tm1c (or floxed) allele by PCR with primers F1 and R2 (wild-type allele “+” 127 bp and floxed allele “fl” 337 bp). Then mice carrying the *Rpa1* tm1c *fl* allele were crossed with *Zp3-Cre* mice (de Vries et al., 2000), where Cre recombinase expression is detected only in oocytes starting from the primary follicles (JAX, #003651), or *Ddx4-Cre* mice (JAX, #006954), where Cre activity is directed to the embryonic germ cells starting at E15 (Gallardo et al., 2007). The resultant tm1d allele (deletion of *Rpa1* Exon 8) was confirmed by PCR with primers F1 and R3 tggttaatgcaaacaggacag (deletion allele “-” 461 bp). Deletion of Exon 8 causes a frame shift in the resulting *Rpa1* mutant transcript. Desired offspring were selected for breeding (no *FLP* allele in breeders) to produce the oocyte-specific conditional knockout (OcKO) females (*Rpa1* ^fl/fl^ with Cre or *Rpa1* ^fl/-^ with Cre) and controls (*Rpa1* ^fl/fl^ without Cre or *Rpa1* ^fl/+^ with Cre).

**Figure 1.**
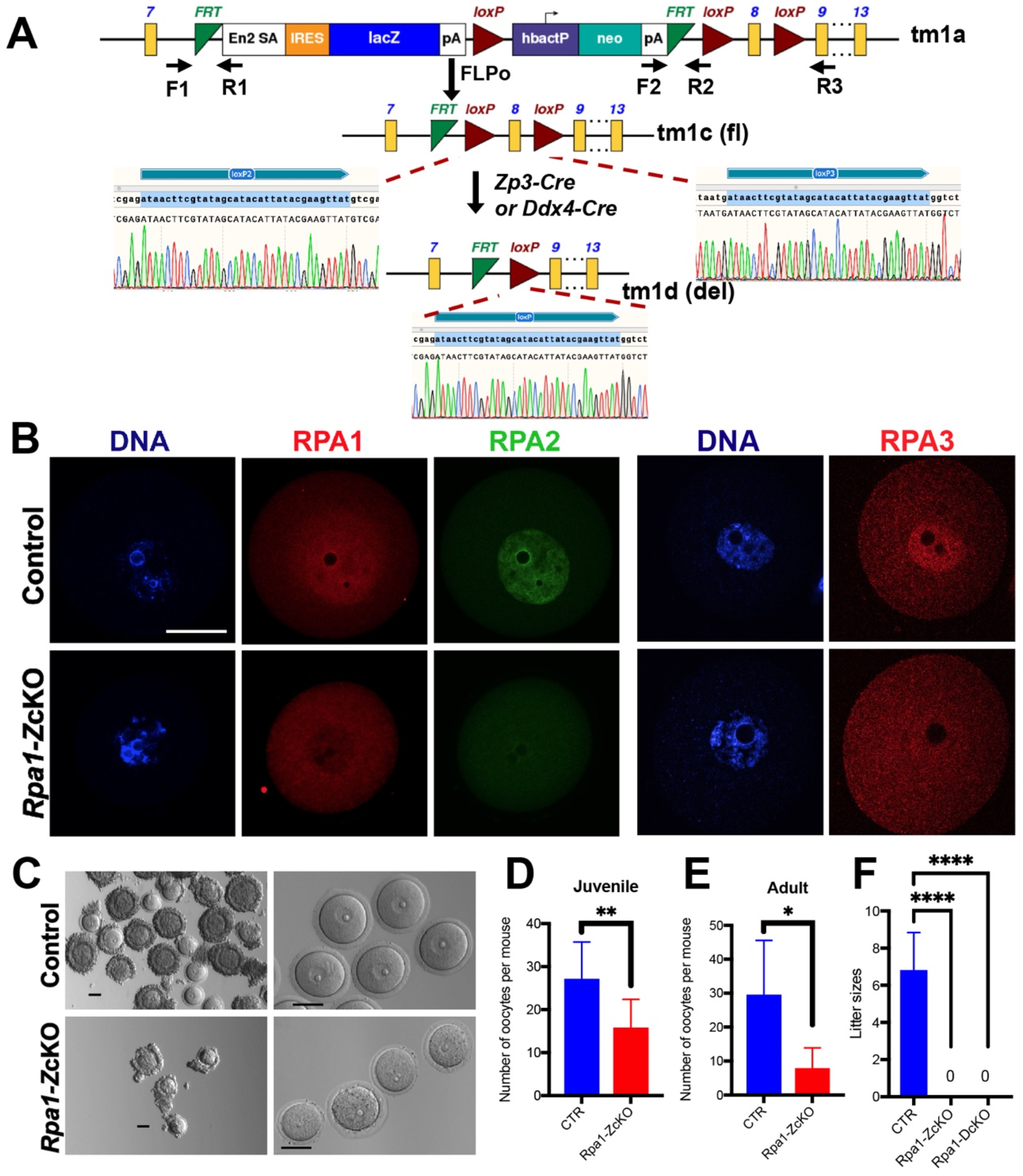
Depletion of RPA1 in oocytes causes loss of the RPA complex and female infertility. (A) Generation of *Rpa1* oocyte-specific conditional knockout (OcKO) mice. (B) Immunofluorescence of RPA1, RPA2, and RPA3 in control (*Rpa1* ^fl/fl^ without Cre or *Rpa1* ^fl/+^ with Cre) and *Rpa1-Z*cKO GV oocytes. Note the non-specific background signals in the cytoplasm of both genotypes in RPA1 and RPA3 images (4- to 5-week-old, n=3 per genotype). (C) Representative images of GV oocytes collected from adult control and *Rpa1-Z*cKO mice, before and after the removal of cumulus cells. For convenient comparison, some scale bars were placed adjacent to the oocytes instead of in the corners. (D, E) Number of GV oocytes obtained from the controls and *Rpa1-Z*cKOs in juvenile (4- to 5-week-old, CTR=9, *Rpa1-Z*cKO=10) and adult mice (8-week-old, n=6 per genotype) after hormone PMSG treatment. (F) Mating test showed the sterility of both *Rpa1-Z*cKO females and *Rpa1-D*cKO females. CTR=6, *Rpa1-Z*cKO=5, *Rpa1-D*cKO=3. Data are presented as mean ± SD, \**P*<0.05, \*\**P*<0.01, \*\*\*\**P*<0.0005. F, forward primer; R, reverse primer. tm, targeted mutation. fl, floxed allele. del, deletion allele. Scale bars: 50 µm.

### Ovary imaging and GV oocyte collection

*Rpa1* OcKO and control female mice were superovulated with 5 IU (juveniles) or 10 IU (adults) pregnant mare serum gonadotropin (PMSG, BioVendor, Asheville) by intraperitoneal injection. After the removal of the ovarian bursa, ovaries were imaged under the same magnification. GV oocytes were collected as described previously (Zhou et al., 2012). Briefly, fully grown GV oocytes were obtained from ovaries 46 hours post hormone treatment by puncturing the antral follicles with a fine needle in M2 medium (MR015D) under a stereomicroscope. For oocyte number counting, all GV oocytes with and without cumulus cells were counted. Then cumulus cells were removed mechanically with a fine glass pipette before the fixation of GV oocytes for future analysis. For oocyte in vitro maturation (IVM), GV oocytes were cultured in CZB medium (MR019D) at 37°C in a humidified atmosphere of 5% CO_2_.

### Immunofluorescence (IF) in oocytes

All steps were carried out at room temperature unless otherwise noted. Oocytes were collected from superovulated 4- to 5-week-old mice. Oocytes were fixed in 4% paraformaldehyde (PFA) for at least 30 min and briefly washed in PBS with 0.3% polyvinylpyrrolidone (PVP), followed by permeabilization with 0.5% Triton X-100 in PBS for 20 min. Oocytes were then placed in blocking solution (PBS with 10% FBS and 0.1% Triton) for 1 h, and incubated with primary antibodies overnight at 4°C. The primary antibodies were diluted in blocking solution with 1:200 dilution unless otherwise noted, including: γH2AX (Ser139), Cell Signaling, 80312; pRb (Ser807/811), Cell Signaling, 8516; Rad51, Cell Signaling, 8875; LAP2 (lamina-associated polypeptide 2, also known as Thymopoietin), Thermo Fisher, PA5-52519; pATM (pS1981, Clone K88-534), BD, 560007 (images shown in this manuscript); pATM (Ser1981, Clone 10H11.E12), Cell Signaling, 4526 (similar results, images not shown); pATR (T1989), Abcam, ab289363; DNA-PK (phospho S2056), Abcam, ab18192; p-P53 (Ser15), Cell Signaling, 9284; acetyl-P53 (Lys379), Cell Signaling, 2570; RPA1, Thermo Fisher, PA5-21976; RPA2, Cell Signaling, 2208; RPA3 (Shi et al., 2019); α-Tubulin, Cell Signaling, 3873; CREST antiserum 1:500, a gift from B.R. Brinkley (Brenner et al., 1981). After three times of wash in PBS with 0.3% PVP and 0.1% Triton, oocytes were incubated with the secondary antibodies (1:500 in blocking solution, Alexa Fluor, Thermo Fisher Scientific) for 1 hour in dark. After three times of wash, DNA was stained with 4,6-diamidino-2-phenylindole (DAPI) before mounting and imaging under Nikon Ts2R Microscope or Leica DM5500B microscope.

### Nuclear surface spread analysis

Surface spreads of oocyte nuclei were performed as described previously (Peters et al., 1997). For nuclear spread analysis, oocytes were retrieved from E18.5 embryos. The primary antibodies used for immunofluorescence were: RPA1, Abcam, ab87272 (1:50); SYCP1, Abcam, Ab15090 (1:200); SYCP2 (Yang et al., 2006) (1:200); SYCP3, Abcam, Ab97672 (1:100). Images of immunolabeled chromosome spreads were captured with an ORCA Flash4.0 digital monochrome camera (Hamamatsu Photonics) on a Leica DM5500B microscope (Leica Microsystems).

### Histological section and staining

Ovaries were fixed in 4% PFA overnight. After wash and dehydration, ovaries were embedded in paraffin, and sectioned at 8-µm intervals and stained with hematoxylin and eosin. For fluorescent detection, after deparaffinization, sections were treated with Tris-EDTA buffer to retrieve antigen and stained with the 1:500 diluted primary antibodies: pATM (pS1981), BD, 560007; pATR (T1989), Abcam, ab289363; DNA-PK (pS2056), Abcam, ab18192.

### Quantification of follicles

Follicle quantification was carried out following the previously described protocol (Myers et al., 2004). Ovaries were sectioned at 8-µm intervals and subjected to hematoxylin and eosin staining. In each mouse, the number of follicles was counted in every fifth section of one ovary, and the sum of oocytes from all counted sections was considered the total number of oocytes per ovary. To prevent duplicate counting, only oocytes with a visible nucleus were included in the count (Zhou et al., 2015). The stages of follicles were identified as follows: primordial follicles (PrF), characterized by a single layer of granulosa cells with flattened nuclei surrounding the oocyte; primary follicles (PF), with a single layer of cuboidal granulosa cells surrounding the oocyte; secondary follicles (SF), featuring more than one layer of cuboidal granulosa cells without a visible antrum; antral follicles (AF), exhibiting antral space within the granulosa cell layers.

### TUNEL assay

Terminal deoxynucleotidyl transferase dUTP nick end labeling (TUNEL) assay was performed using the In Situ Cell Death Detection Kit (11684795910, Roche) according to the manufacturer’s protocol. Oocytes were washed and labeled with TUNEL reaction mixture at 37°C for 30 min (protected from light). For tissue sections, fluorescent detection was performed after deparaffinization. Sections were treated with 0.1 M citrate buffer, pH 6 before the TUNEL reaction according to the protocol.

### RNA-seq analysis

Fully grown GV oocytes were collected from superovulated 4- to 5-week-old mice. Three biological replicates were prepared for each genotype and each sample contained around 20 GV oocytes without any cumulus cells. RNA-seq library construction, sequencing, and data analysis were undertaken by Azenta Life Sciences (NJ, USA) under ultra-low input RNA-seq workflow. Sequence reads were trimmed to remove possible adapter sequences and nucleotides with poor quality using Trimmomatic v.0.36. The trimmed reads were mapped to the *Mus musculus* GRCm38 reference genome available on ENSEMBL using the STAR aligner v.2.5.2b. Unique gene hit counts were calculated and only unique reads that fell within exon regions were counted, which produced a total of 379,525,638 reads and 113,857 Mbases yield obtained from 6 samples (Control=3, OcKO=3) with 92.62% bases >= Q30. DESeq2 and Wald test were used to generate p-values and log2 fold changes. Genes with an adjusted p-value < 0.05 and absolute log2 fold change > 1 were called as differentially expressed genes (DEGs). Gene Ontology (GO) analysis was performed on the statistically significant set of genes by implementing the software GeneSCF v.1.1-p2, clustering the set of genes based on their biological processes and statistical significance.

### Fertility testing

Six to seven-week-old control and *Rpa1* OcKO female mice were mated continuously with 8- to 10-week-old wild-type C57BL/6N fertile males for three months. The number of pups on the first day after parturition was counted as the litter size.

### Image analysis

Each batch of images containing both control and OcKO oocytes were acquired under identical capture settings, and relative intensities were measured on the raw images using Fiji software (Schindelin et al., 2012). During intensity analysis, signals originating from cumulus cells were excluded. The control group served as the standard, and the relative intensity of the OcKO group was compared to the control group.

### Statistical analysis

Statistical analyses were conducted using GraphPad Prism software version 8.0. An unpaired two-tailed Student’s t-test was employed to compare the data between control and OcKO mice, with a significance level set at P<0.05. The results are presented as mean ± standard deviation (SD). Each experiment was performed at least three times to ensure reproducibility. The number of samples utilized in each experiment is indicated in the figure legends.

## Results

### Loss of RPA1 in postnatal oocytes causes inactivation of the RPA complex and female infertility

*Rpa1* oocyte-specific conditional knockout (OcKO) female mice were generated as described in the Methods section (Fig. 1A). The deletion of exon 8 causes a frame shift in the resulting mutant transcript. To validate this, we performed immunofluorescence (IF) analysis using an RPA1-specific antibody that has been previously validated in both mouse cell lines (Flach et al., 2014) and mouse oocytes (Yueh et al., 2021). RPA1 was present in the nucleus of wild type GV oocytes, indicating that RPA may play an additional role in oocytes after meiotic recombination. The RPA1 signal was absent in the nucleus of GV oocytes collected from the *Zp3-*Cre-mediated *Rpa1* OcKOs (hereafter referred to as *Rpa1-Z*cKO, Fig. 1B), suggesting efficient depletion of RPA1 after *Zp3*-Cre-mediated deletion. Since RPA is a heterotrimer composed of RPA1, RPA2, and RPA3, we further examined the other RPA subunits using specific antibodies (Shi et al., 2019). Our results demonstrate that loss of RPA1, the largest subunit, leads to the loss of the entire RPA complex in oocytes, as evidenced by the disappearance of RPA2 and RPA3 (Fig. 1B). This result in oocytes is consistent with the loss of the RPA complex in RPA1-deficient spermatocytes (Shi et al., 2019). Moreover, loss of RPA1 resulted in a decrease in the number and size of GV oocytes (Fig. 1C), and this decrease in oocyte number appeared to be age-dependent, with a more severe phenotype observed in adult *Rpa1-Z*cKO mice compared to the juveniles (Fig. 1D, E). Finally, mating tests confirmed the sterility of *Rpa1-Z*cKO females, which was also observed using another *Cre* driver, *Ddx4-Cre* (*Rpa1-D*cKO, Fig. 1F). These results demonstrate that RPA plays an essential role in oocytes after meiotic recombination.

### Loss of RPA1 in oocytes impairs ovarian folliculogenesis

We proceeded to investigate the underlying cause of sterility in *Rpa1* OcKO females. At approximately 3.5 weeks of age, the size of ovaries from *Rpa1-Z*cKO females was comparable to that of the control group. However, by 5 weeks of age, the OcKO females exhibited smaller ovarian size, and no further growth was observed beyond 10 weeks of age (Fig. 2A). Furthermore, even after hormone PMSG treatment, very few large follicles could be observed in the OcKO ovaries at 10 and 16 weeks of age (Fig. 2A). To further confirm these observations, histological ovarian sections were analyzed, and follicle quantification was performed across various age groups, ranging from 2 months to 8 months (Fig. 2B). At 2 months of age, the number of antral follicles was significantly decreased in the OcKO females (Fig. 2C). Starting at 4 months of age, both secondary and antral follicles exhibited a significant decrease in numbers (Fig. 2D). These findings indicate that the severity of oocyte depletion increases with age in OcKO females. Although the numbers of primordial and primary follicles were comparable between control and OcKO females, the total number of follicles decreased in the *Rpa1-Z*cKO ovaries due to the severe reduction of secondary and antral follicles (Fig. 2D-F). As a result, the OcKO ovaries were smaller (Fig. 2A). Taken together, postnatal inactivation of RPA causes a substantial loss of secondary and antral follicles.

**Figure 2.**
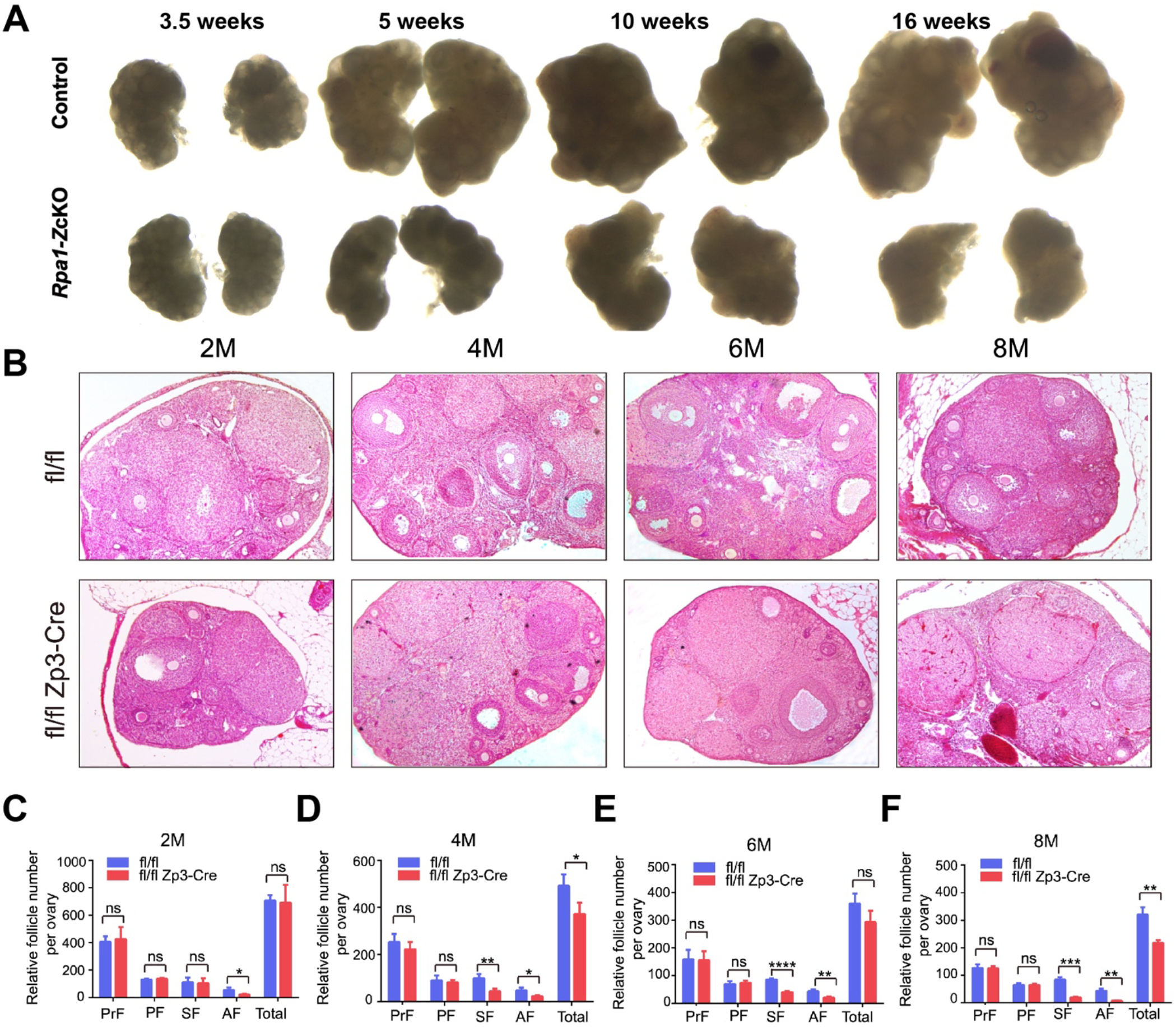
Loss of RPA1 in oocytes impairs ovarian folliculogenesis. (A) Representative images of ovaries from mice at different ages (under the same magnification). Ovaries were dissected after PMSG treatment (n=3 per genotype per age). (B) Histological analysis of ovaries at 2, 4, 6, and 8 months in control and OcKO mice without hormone treatment (n=3 per genotype per age). (C-F) Follicle quantification in control and OcKO females at different ages. PrF, primordial follicles; PF, primary follicles; SF, secondary follicle; AF, antral follicle. M, month. Data are presented as mean ± SD. \**P*<0.05, \*\**P*<0.01, \*\*\**P*<0.001, \*\*\*\**P*<0.0005, ns, not significant.

### Occurrence of DNA breaks in granulosa cells in *Rpa1*-*Z*cKO antral follicles

To investigate the underlying cause of the substantial loss of secondary and antral follicles in *Rpa1*-*Z*cKO ovaries, immunofluorescence analysis was performed on ovarian sections. Interestingly, although minor DNA strand breaks detected by TUNEL were shown in granulosa cells of the control ovaries, massive TUNEL signals were observed specifically in the granulosa cells of the antral follicles, but not other follicles, in the *Rpa1-Z*cKO ovaries (Fig. 3A, B). Given that the Cre-recombinase activity under *Zp3-Cre* is specific to oocytes at the primary follicle stage (de Vries et al., 2000; Sun et al., 2008), the observed severe DNA breaks in the granulosa cells at the antral follicle stage was not likely due to leaked Cre activity. Additionally, the granulosa cells showing DNA breaks were not the closest cells to the oocytes, ruling out the possibility of diffusion of Cre from the oocyte to its surrounding cells. To further investigate the DNA damage response, we examined the levels of activated ATM, ATR, and DNA-PK, which are key kinases involved in the DNA damage response. Although there was a slight increase in the activated kinases in *Rpa1*-*Z*cKO ovaries, the difference was not statistically significant (Fig. 3C-G).

**Figure 3.**
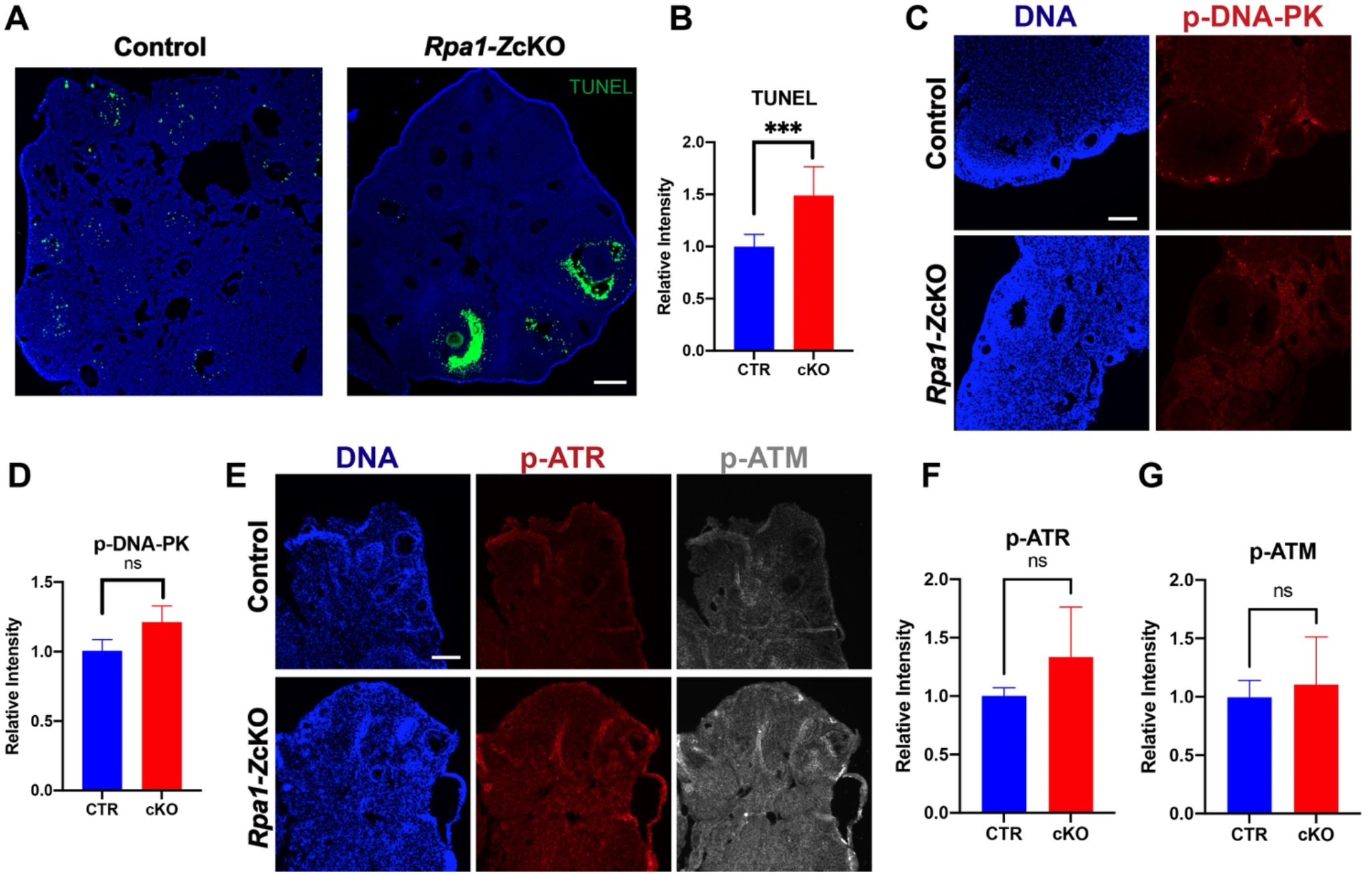
Occurrence of DNA breaks in granulosa cells in *Rpa1*-*Z*cKO antral follicles. (A, B) Immunostaining and quantification of TUNEL signal in ovarian sections of control and *Rpa1-Zp3* OcKO females (4- to 5-week-old, n=3 per genotype). (C, D) Immunostaining and quantification of p-DNA-PK (phospho S2056) in ovarian sections of control and *Rpa1-Zp3* OcKO females (5- to 6-week-old, n=3 per genotype). (E, F, and G) Immunostaining and quantifications of p-ATR (phospho T1989) and p-ATM (phospho S1981) in ovarian sections of control and *Rpa1-Zp3* OcKO females (6-week-old, n=5 per genotype). Data are presented as mean ± SD. \*\*\**P*<0.001, ns, not significant. Scale bars: 100 µm.

### Non-DNA break mediated DNA damage exists in *Rpa1*-*Z*cKO oocytes

We further investigated the phenotypes of *Rpa1*-*Z*cKO oocytes. Given that RPA1 plays a crucial role in maintaining genome integrity through its strong interaction with LAP2 (lamina-associated polypeptide 2, also known as Thymopoietin) (Bao et al., 2022), we analyzed the levels of LAP2 and phosphorylated histone H2AX at Ser139 (γH2AX), a well-known marker of DNA damage (Fig. 4A). We found a significant increase in both LAP2 and γH2AX in *Rpa1*-OcKO oocytes, indicating a potential alteration in LAP2-mediated nuclear membrane-chromatin attachment (Gant et al., 1999) due to the loss of RPA1. Consequently, this alteration may trigger the recruitment of repair factors, as evidenced by the dramatic accumulation of γH2AX in RPA1-deficient oocytes (Fig. 4B, C). To assess if the accumulation of γH2AX resulted from DSBs, we examined RAD51 recombinase, a protein involved in DSB repair in coordination with the RPA complex (Rinaldi et al., 2017; Stauffer and Chazin, 2004). However, no significant change in RAD51 levels was observed, indicating either the absence of DSBs or an inactive DSB repair pathway in *Rpa1*-OcKO oocytes (Fig. 4D, E). Additionally, the TUNEL assay, capable of detecting both SSBs and DSBs, did not detect apparent DNA strand breaks in *Rpa1*-OcKO oocytes (Fig. S1). In conclusion, *Rpa1-Z*cKO oocytes displayed severe DNA damage, which is not attributed to DNA breaks.

**Figure 4.**
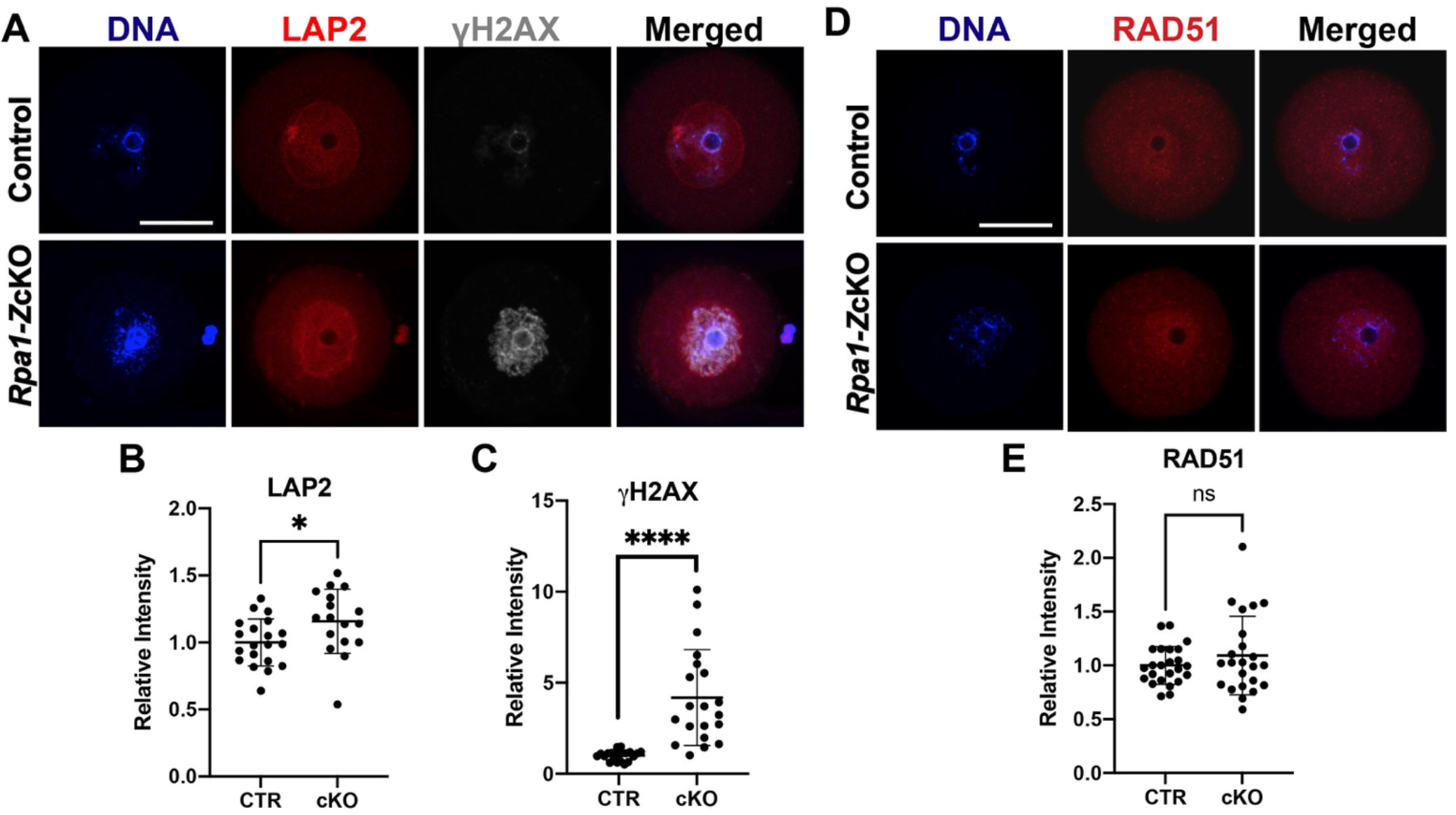
Non-DNA break-mediated DNA damage exists in *Rpa1*-*Z*cKO oocytes. (A) Immunostaining of LAP2 and γH2AX in control and *Rpa1-Z*cKO oocytes. (B, C) Quantifications of LAP2 (CTR=20, OcKO=17) and γH2AX (CTR=21, OcKO=20). Three mice per genotype, 4-5 weeks of age, were used in the experiment. (D, E) Immunostaining and quantification of RAD51 in control and *Rpa1-Z*cKO oocytes (CTR=24, OcKO=23). Three mice per genotype, 4-5 weeks of age, were used in the experiment. Signals originating from cumulus cells were excluded during the intensity analysis. Data are presented as mean ± SD, \**P*<0.05, \*\*\*\**P*<0.0005, ns, not significant. Scale bars: 50 µm.

### P53 and Rb exhibit significant phosphorylation in *Rpa1*-*Z*cKO oocytes

In order to assess the impact of RPA loss in fully grown GV oocytes, we focused on p53 (known as TRP53 in mice), since p63 expression is absent in antral follicles (Livera et al., 2008; Suh et al., 2006). Notably, *Rpa1*-OcKO oocytes exhibited a significant increase in phosphorylation of p53 at Ser15, indicating activation of the p53 pathway in the absence of the RPA complex (Fig. 5A, B). However, there was no noticeable change in the acetylation status of p53 at Lys379 (Fig. 5C, D), suggesting that the loss of RPA complex may not directly influence p53 acetylation or that alternative post-translational modifications of p53 are involved. We also examined the retinoblastoma tumor suppressor protein Rb, which plays a key role in DNA damage response (Wang et al., 2001). We observed a significant increase in phospho-Rb (Ser807/811) levels in *Rpa1*-OcKO oocytes (Fig. 5E, F), suggesting that phosphorylation-mediated inactivation of Rb, a phenomenon commonly observed in tumors or cells with aberrant cell cycles (Wang et al., 2001), is induced in *Rpa1*-OcKO oocytes. These results show that the DNA damage response pathways are activated in RPA-deficient oocytes.

**Figure 5.**
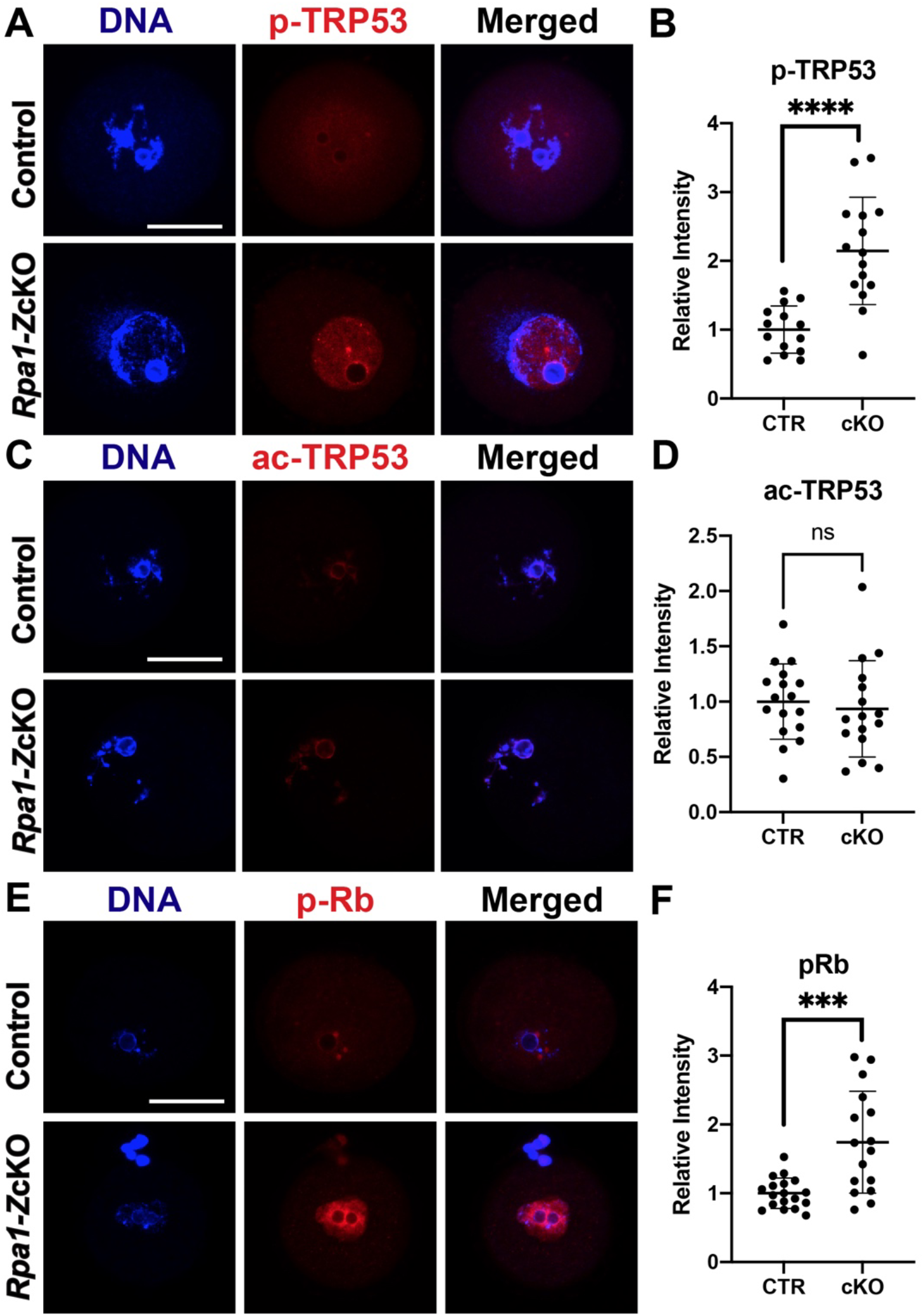
P53 and Rb show pronounced phosphorylation in *Rpa1*-*Z*cKO oocytes. (A, B) Immunostaining and quantification of p-TRP53 (Ser15) in control (n=14) and *Rpa1-Z*cKO oocytes (n=15). (C, D) Immunostaining and quantification of ac-TRP53 (Lys379) in control (n=17) and *Rpa1-Z*cKO oocytes (n=16). (E, F) Immunostaining and quantification of p-Rb (Ser807/811) in control (n=18) and *Rpa1-Z*cKO oocytes (n=16). Three mice per genotype, 4-5 weeks of age, were used in each experiment. Signals originating from cumulus cells were excluded during the intensity analysis. Data are presented as mean ± SD, \*\*\**P*<0.001, \*\*\*\**P*<0.0005, ns, not significant. Scale bars: 50 µm.

### DNA damage response kinases ATM, ATR, and DNA-PK are activated in *Rpa1*-*Z*cKO oocytes

To investigate whether p53 activation occurs in *Rpa1*-*Z*cKO oocytes as a response to DNA damage, we assessed the activity of key kinases involved in DNA damage detection and response, as well as initiation of signaling cascades in the DNA repair processes, namely ATM, ATR, and DNA-PK. In control oocytes, the levels of phosphorylated ATM (p-ATM, S1981) and phosphorylated ATR (p-ATR, T1989) were found to be low. However, in *Rpa1*-OcKO oocytes, both p-ATM and p-ATR exhibited significant increases (Fig. 6A-C). Similarly, while p-DNA-PK (S2056) was barely detectable in control oocytes, its level was significantly elevated in *Rpa1*-OcKO oocytes (Fig. 6D, E). These findings show that DNA damage response and repair processes are activated in *Rpa1*-*Z*cKO oocytes. To profile transcriptomes in fully grown GV oocytes, we performed RNA-seq analysis (6 samples, Control=3, OcKO=3) (Table S1). Among the 641 differentially expressed genes (DEGs), 479 were upregulated, and 162 were downregulated (Fig. 6F and Table S2). Notably, among the top 30 DEGs (Fig. 6G), all were upregulated except *Rpa1*. Among those upregulated DEGs, many are well-known participants in DNA damage response, G2/M DNA damage checkpoint regulation, and apoptosis, including *Pmaip1*, *Btg2*, *Mdm2*, *Ccng1*, and *Eda2r* (Fig. 6F, G). In summary, these results indicate that loss of RPA leads to a pronounced canonical DNA damage response in GV oocytes.

**Figure 6.**
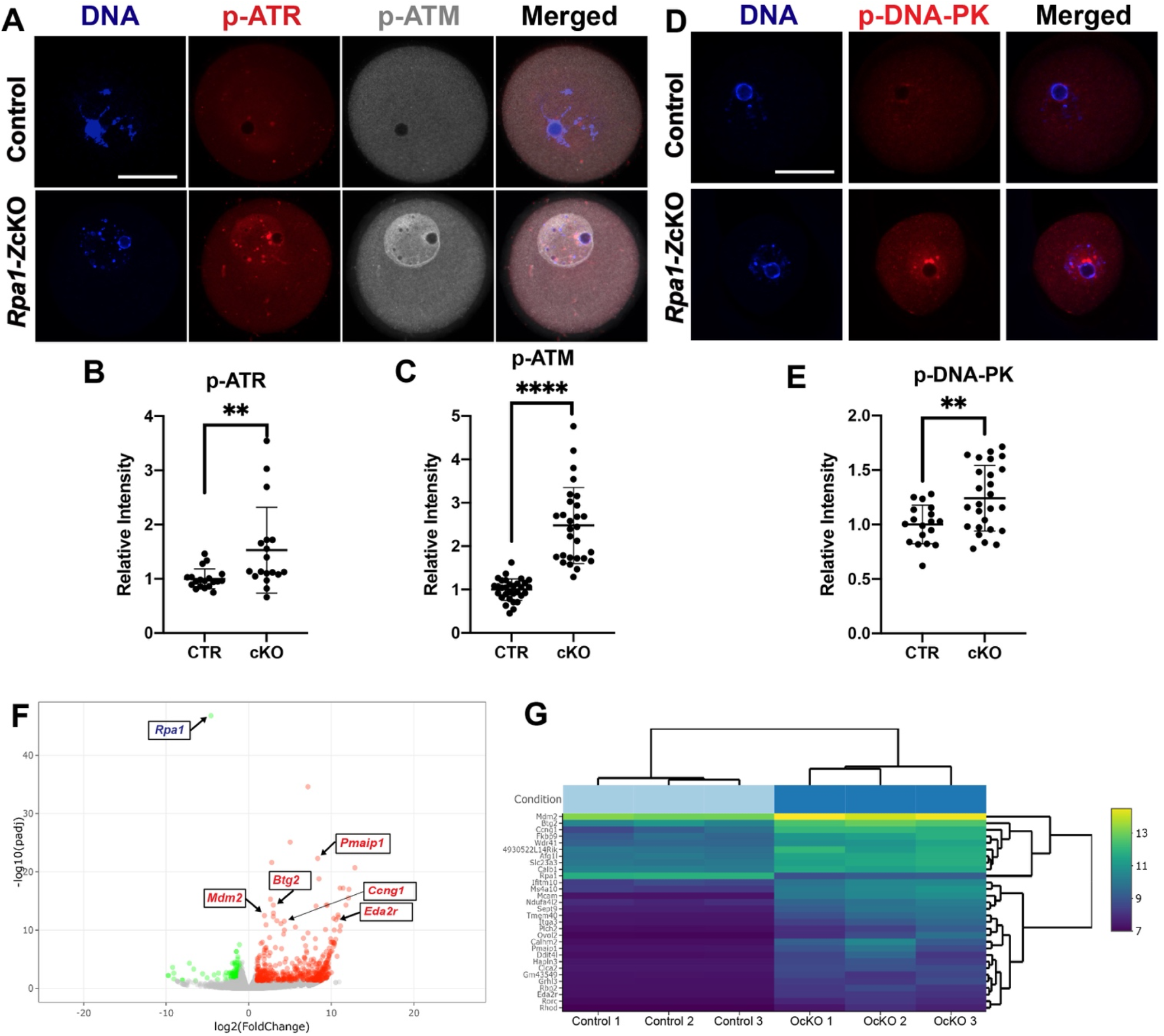
Activation of DNA damage response kinases and transcriptomic changes in *Rpa1*-*Z*cKO oocytes. (A) Immunostaining of p-ATR (T1989) and p-ATM (S1981) in control and *Rpa1*-OcKO oocytes. (B, C) Quantifications of p-ATR (CTR=19, OcKO =18) and p-ATM (CTR=31, OcKO =28). (D, E) Immunostaining and quantification of p-DNA-PK (S2056) in control (n=18) and OcKO oocytes (n=26). (F) RNA-seq analysis revealed 641 differentially expressed genes (DEGs) including 479 upregulated (red dots) and 162 downregulated (green dots) in *Rpa1*-*Z*cKO oocytes. (G) Heatmap of top 30 DEGs, many of which are well-known players in DNA damage response, G2/M DNA damage checkpoint regulation, and apoptosis, including *Pmaip1*, *Btg2*, *Mdm2*, *Ccng1*, and *Eda2r* (highlighted in F). Three mice per genotype, 4-5 weeks of age, were used in each experiment. Data are presented as mean ± SD, \*\**P*<0.01, \*\*\*\**P*<0.0005. Scale bars: 50 µm.

### Loss of RPA in oocytes disrupts chromosome alignment during meiotic divisions

Previously, RPA was shown to be essential for meiotic recombination in mouse spermatogenesis (Shi et al., 2019). To investigate its role in meiotic recombination during oogenesis, we inactivated *Rpa1* using the *Ddx4-Cre*, which begins to express in embryonic germ cells at E15 (Gallardo et al., 2007). Immunofluorescence analysis of nuclear spreads revealed the presence of RPA1 foci on the chromosomes of *Rpa1-Ddx4-*Cre OcKO oocytes at the pachytene stage from E18.5 embryos, indicating that RPA1 was not depleted in the oocytes (Fig. 7A). Furthermore, immunofluorescence of synaptonemal complex proteins showed normal chromosomal synapsis in control and *Rpa1-Ddx4* OcKO oocytes (Fig. 7A, B). The persistence of RPA1 in E18.5 oocytes following *Ddx4-*Cre-mediated deletion is expected. *Ddx4*-Cre begins expression at E15 and thus existing *Rpa1* transcripts and RPA1 protein have not been degraded at E18.5. Therefore, this caveat prevents the assessment of the role of RPA in chromosomal synapsis and meiotic recombination during oogenesis. To allow sufficient time for existing *Rpa1* mRNA and protein degradation, oocytes were collected from P28 mice for in vitro maturation (IVM) and subsequent chromosome alignment analysis. Our results revealed that the majority of *Rpa1-Ddx4* OcKO oocytes exhibited chromosome misalignment at both the metaphase I stage (MI) (Fig. 7C, D) and the metaphase II stage (MII) (Fig. 7E, F), indicating that the DNA damage resulting from RPA loss in GV oocytes persisted and led to chromosome misalignment during meiotic divisions.

**Figure 7.**
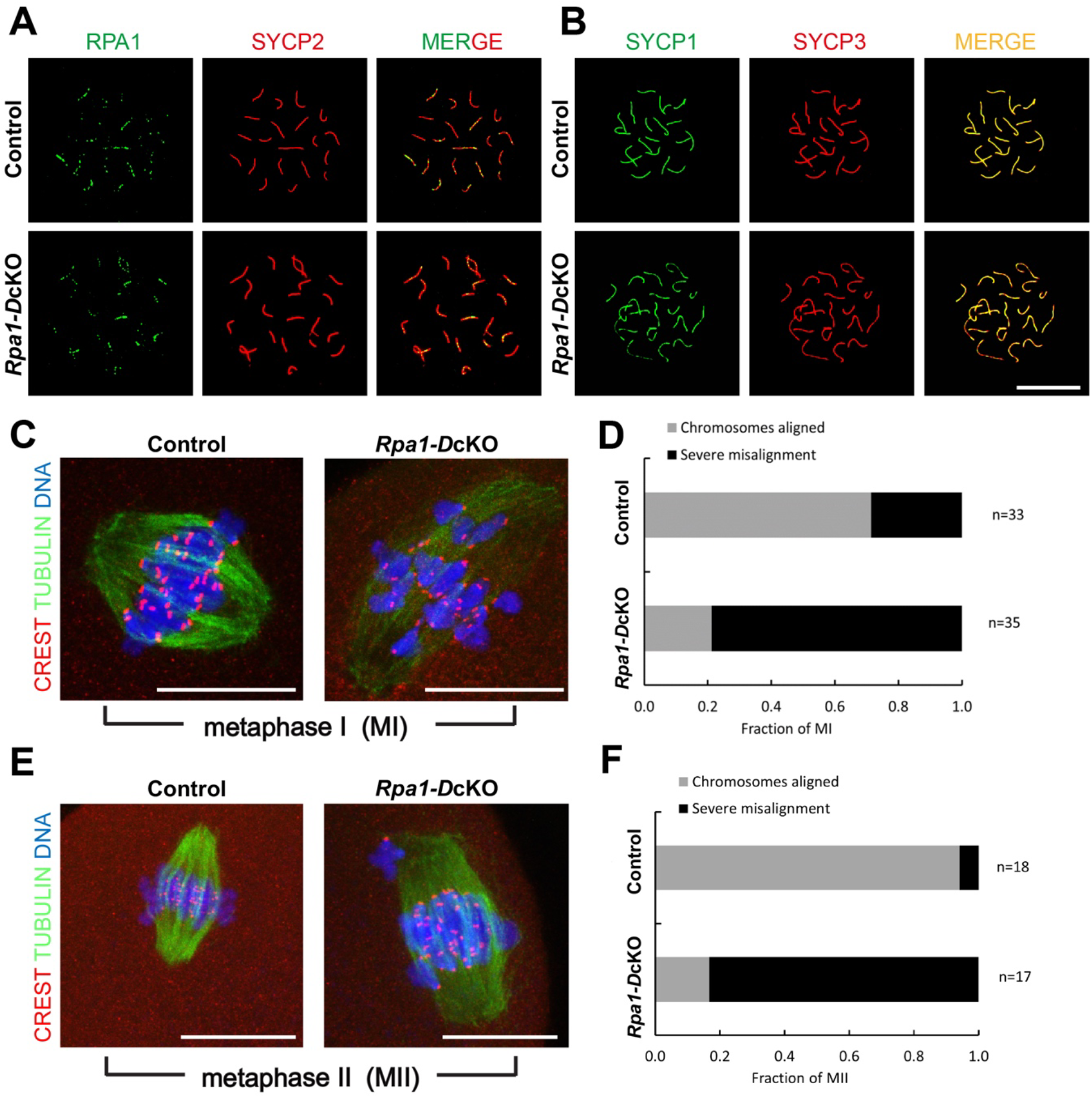
Analysis of meiotic progression and chromosome alignment in *Rpa1-Ddx4-*Cre OcKO oocytes. (A) Immunofluorescence of spread nuclei of pachytene oocytes collected from E18.5 control and *Rpa1-Ddx4*-Cre OcKO embryos with anti-RPA1 and anti-SYCP2 antibodies. (B) Immunofluorescence of synaptonemal complex proteins SYCP1 and SYCP3 in pachytene oocytes from E18.5 embryos. (C, D) Chromosome alignment of control and *Rpa1-Ddx4*-Cre OcKO oocytes at metaphase I stage (MI) collected from P28 mice. (E, F) Chromosome alignment at metaphase II stage (MII) with staining on tubulin (green), kinetochores (CREST, red) and DNA (DAPI, blue). Four mice per genotype were used in each experiment. N, number of oocytes. Scale bars: 25 µm.

## Discussion

Although RPA has been studied in yeast and cell lines, its role in mammalian development remains largely unexplored due to the early embryonic lethality resulting from *Rpa1* mutation (Wang et al., 2005) and null allele (Wang et al., 2015) in mice. In this study, we utilized two *Cre* drivers and conditional KO strategy specifically in oocytes to investigate the consequences of RPA1 loss. We find that depletion of RPA1, the largest subunit of the RPA complex, results in the loss of the entire RPA complex in oocytes, consistent with the previous observation in male germ cell-specific *Rpa1* conditional knockout mice (Shi et al., 2019). Furthermore, the absence of RPA2 and RPA3 proteins in *Rpa1*-OcKO oocytes is likely due to disruption of protein-protein interactions rather than transcriptional inhibition, because RNA-seq data (Table S1) show no changes in their transcript levels.

RPA is known for its high affinity binding to ssDNA, effectively coating and protecting it against degradation and secondary structure formation (Deng et al., 2015). Previous research has primarily focused on RPA’s role at DNA replication forks, where long stretches of ssDNA are present during replication stress or at DNA damage sites (Dodson et al., 2004; Zou et al., 2006). However, in our study, we focused on GV oocytes, which have already completed premeiotic DNA replication before entering prophase of meiosis I. This rules out the possibility of DNA damage accumulation during replication (S phase) in the absence of RPA. Instead, growing GV oocytes exhibit robust transcription activity, resulting in the accumulation of transcripts and proteins, as well as an increased volume around 150-fold over a 3-week period (Svoboda et al., 2015; Tora and Vincent, 2021). Based on this premise, we postulate that the severe DNA damage responses observed in *Rpa1*-OcKO oocytes are likely attributed to transcription-associated DNA damage, which can arise due to collisions between transcription machineries (DNA into RNA), DNA topological stress, or the formation of three-stranded DNA/RNA hybrids (also known as R-loops) during meiosis (Fujiwara et al., 2022) when RPA is absent. This explanation is supported by both the TUNEL assay and RAD51 recombinase immunofluorescence results, which showed the absence of DNA breaks and an inactive DSB repair pathway in *Rpa1*-OcKO oocytes, respectively. The activation of p-ATM, p-ATR, p-DNA-PK, p-p53, and γH2AX further supports the occurrence of DNA damage and the activation of DNA damage response pathways in *Rpa1*-OcKO oocytes. However, elucidating the precise mechanisms by which the loss of the RPA complex in oocytes leads to the activation of these pathways and the pronounced and sustained DNA damage necessitates further investigation.

Our study provides a unique mouse model to investigate the G2/M checkpoint in GV oocytes. Previous studies have often relied on chemical induced DSBs to study the DNA damage checkpoint in G2/prophase oocytes. It is generally accepted that fully grown GV oocytes have a weakened G2/M checkpoint due to limited activation of the ATM kinase (Marangos and Carroll, 2012). However, in our study, we observed robust activation of ATM in *Rpa1*-OcKO oocytes (confirmed by two different commercial antibodies), suggesting that non-DSB-mediated DNA damage might be more efficient than DSBs in activating ATM in G2/prophase oocytes. Nevertheless, despite the activation of ATM, these *Rpa1*-OcKO oocytes still progressed to the MI stage (Fig. 7), indicating a weakened G2/M checkpoint in oocytes, consistent with previous studies (Marangos and Carroll, 2012; Subramanian et al., 2020). Interestingly, *Rpa1*-OcKO oocytes were able to reach the MII stage, which differs from DNA damage induced by radiation, ultraviolet B, or chemicals that typically result in MI arrest due to the spindle assembly checkpoint (SAC) response (Collins and Jones, 2016; Collins et al., 2015; Pailas et al., 2022). This suggests that the DNA damage resulting from RPA loss in oocytes is distinct from other types of DNA damage reported previously. Notably, while *Rpa1*-OcKO oocytes were able to progress to MI and MII, we observed severe chromosome misalignment in these oocytes, indicating that DNA damage carried over from the GV stage affects chromosome alignment during subsequent meiotic divisions. Although the precise mechanisms underlying this phenomenon require further investigation, our RNA-seq Gene Ontology (GO) analysis revealed that actin cytoskeleton organization was most significantly altered in response to the loss of RPA in oocytes (Table S3). Previous studies have demonstrated that aberrant actin can lead to increased chromosome misalignment during MI and MII (Blengini and Schindler, 2022; Duan and Sun, 2019; Mogessie and Schuh, 2017; Roeles and Tsiavaliaris, 2019). Actin also interacts with other nuclear proteins and actively participates in various DNA-related processes, including DNA repair (Hurst et al., 2019), which may explain the observed increase of LAP2 in *Rpa1*-OcKO oocytes.

The age-dependent progressive oocyte depletion phenotype in *Rpa1*-OcKO females is intriguing. As these females age, the phenotypes worsen, including a decline in oocyte numbers and the loss of secondary and antral follicles under the *Zp3-Cre* driver (Fig. 1 and Fig. 2). Similarly, when the earlier driver *Ddx4-Cre* was used, females exhibited an even earlier and more severe phenotype, with the loss of almost all follicles, including primordial and primary follicles, as early as 4-6 weeks of age (Fig. S2). These findings suggest that it takes some time for the pre-existing *Rpa1* mRNA and protein to degrade fully and for the phenotypes to manifest after Cre-mediated deletion. This explains the detection of RPA1 on E18.5 oocyte chromosomes after *Ddx4-*Cre-mediated deletion beginning at E15. The recently developed *Stra8^P2Acre^* (Ahmed et al., 2023), which initiates expression in female germ cells at E13, may be useful for investigating this aspect. Importantly, our *Rpa1*-OcKO models support the theory that the oocyte plays a crucial role in determining the overall rate of follicular development (Eppig, 2001; Eppig et al., 2002; Reddy et al., 2008). This is evident from the strong TUNEL signals observed in granulosa cells and the loss of follicles following depletion of RPA in oocytes. We hypothesize that GV oocytes carrying severe DNA damage due to RPA loss fail to drive the growth and/or differentiation of certain granulosa cells through paracrine factors, but the specific mechanisms require further investigation.

Our study provides valuable insights into the role of the RPA complex during mammalian oogenesis. We have demonstrated that RPA is crucial for DNA damage repair, and its loss leads to extensive and persistent DNA damage responses, as evidenced by the activation of key signaling molecules including p-ATM, p-ATR, p-DNA-PK, p-p53, and γH2AX. Additionally, the loss of RPA in oocytes triggers an increase in chromatin-binding nuclear protein LAP2 as well as expression of genes regulating G2/M DNA damage checkpoint and apoptosis, such as *Pmaip1*, *Btg2*, *Mdm2*, *Ccng1*, and *Eda2r*. During MI and MII meiotic divisions, severe chromosome misalignment also occurs in RPA-deficient oocytes. Furthermore, our oocyte-specific conditional knockout models reveal the significant role of mammalian oocytes in orchestrating ovarian follicle development. Specifically, the loss of RPA in oocytes leads to programmed cell death in granulosa cells, resulting in the depletion of growing follicles and a decline in oocyte numbers, ultimately leading to female infertility. These findings shed light on the intricate mechanisms underlying DNA damage responses in G2/prophase oocytes after meiotic recombination and highlight the critical role of RPA in preserving oocyte quality and female reproductive health.

## Supplementary information

**Fig. S1.**
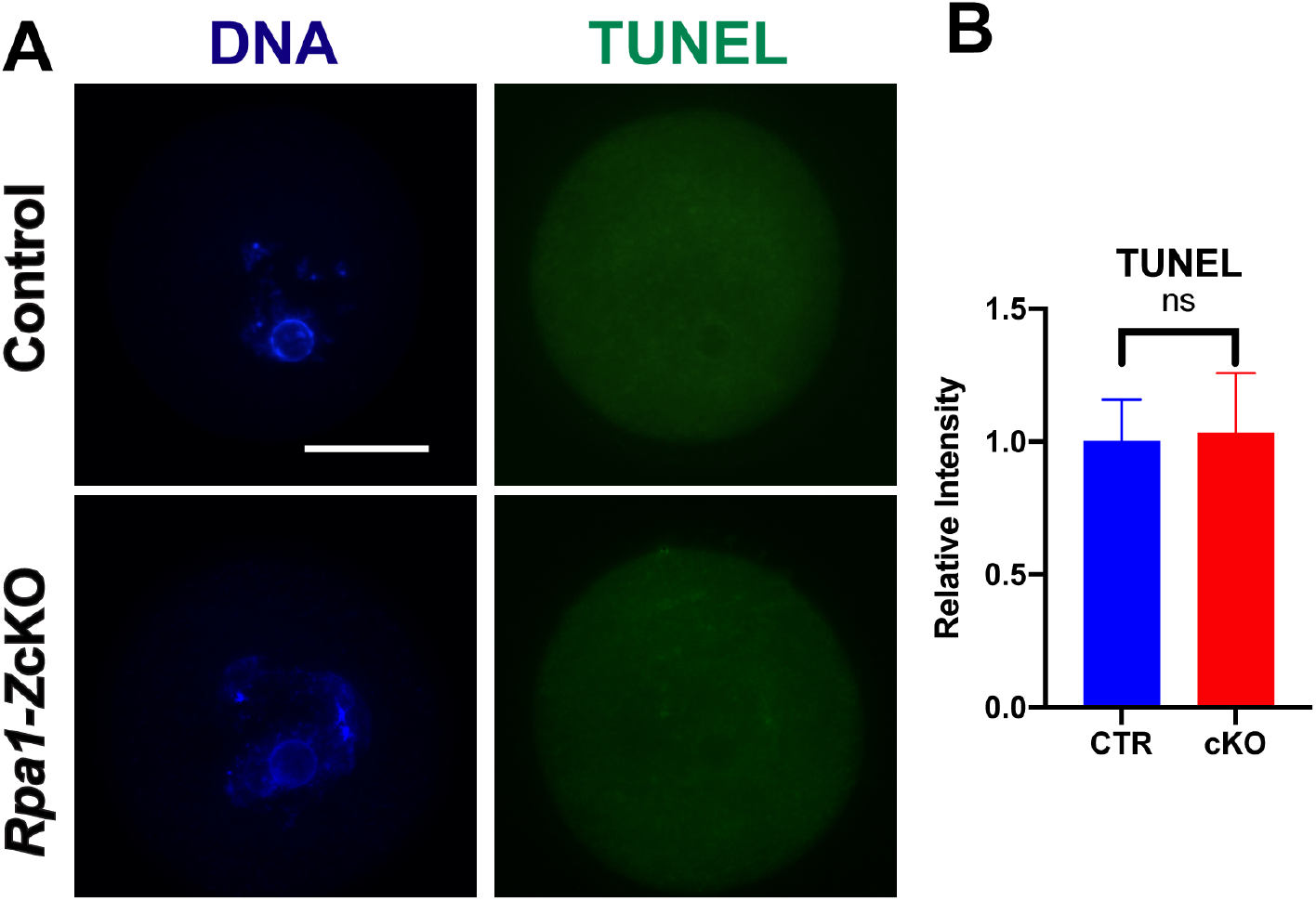
No obvious DNA breaks were detected by TUNEL assay in *Rpa1*-*Z*cKO oocytes. (A) Representative images of TUNEL assay in control and *Rpa1*-OcKO GV oocytes. (B) Quantification of TUNEL signal in control (n=29) and *Rpa1*-OcKO (n=34) oocytes. Four mice per genotype, 4-5 weeks of age, were used in the experiment. Data are presented as mean ± SD. ns, not significant. Scale bar: 50 µm.

**Fig. S2.**
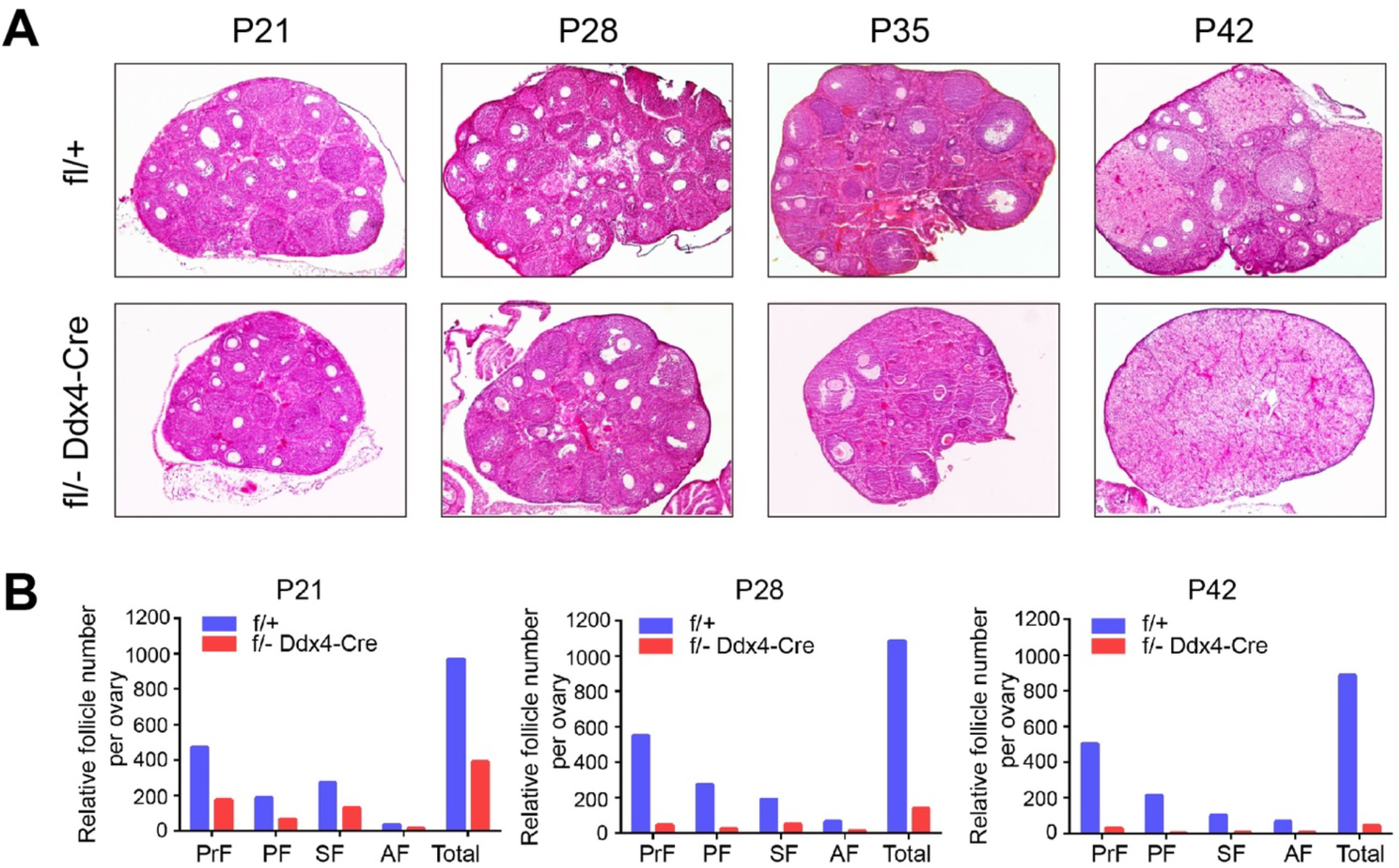
Earlier and more severe loss of follicles, including primordial and primary follicles, were detected under the earlier driver *Ddx4-Cre*. (A) Histological analysis of ovaries at 3, 4, 5, and 6 weeks in control and OcKO females without hormone treatment. (B) Follicle quantification in control and OcKO females at different ages. PrF, primordial follicles; PF, primary follicles; SF, secondary follicle; AF, antral follicle. P, postnatal day. Four mice per genotype (1 per each age) were used in the experiment.

Table S1-*Rpa1* OcKO-RNAseq full list

Table S2-*Rpa1* OcKO-Significant-DEGs

Table S3-*Rpa1* OcKO-GO_analysis

## Acknowledgements

The authors thank the Knockout Mouse Project (KOMP) and Mutant Mouse Resource & Research Centers (MMRRC, University of California, Davis) for providing *Rpa1* tm1a allele (originating from Stephen Murray, The Jackson Laboratory). We thank the Animal Models Core Facility at the Institute for Applied Life Sciences (IALS) of University of Massachusetts-Amherst for performing the IVF and embryo transfer work. We thank Rafael Fissore (UMASS Amherst) for sharing the *Zp3-Cre* mouse line. We are grateful to Tieqi Sun and Angela Houh of Cui laboratory for assistance in genotyping.

## Data availability

All the data were provided in the manuscript and the supplementary information.

